# The assembly of Y chromosome reveals amplification of genes regulating male fertility in *Bactrocera dorsalis*

**DOI:** 10.1101/2024.08.01.606120

**Authors:** Shuangxiong Wu, Jiahong Wu, Quan Lei, Donghai He, Xinrui Jiang, Chao Ye, Dong Wei, Jinjun Wang, Luohao Xu, Hongbo Jiang

## Abstract

**BACKGROUND:** The oriental fruit fly *Bactrocera dorsalis* is an invasive pest causing considerable ecological and economic damage worldwide. The Y chromosome is an important target for the sterile insect technique (SIT) but its sequence and structure has been poorly explored.

**RESULTS:** We assembled the genome of *B. dorsalis* at the chromosome level with a total size of 562.6 Mb. The assembly includes a ∼7.6 Mb Y chromosome sequence, being the first reported Y chromosome in Tephritidae. The assembled Y chromosome is highly repetitive, and contains 61 genes, including 9 multi-copy genes. We surprisingly found that the M factor (*MoY*) in Tephritidae has multiple copies, verified by our droplet digital PCR (ddPCR) analysis. Besides, we identified 32 copies of *gyf-like on the Y chromosome* (*gyfY*) that were highly expressed in testis. RNAi targeting the *gyfY* resulted in depletion of live sperms, suggesting that the amplification of *gyfY* is essential for male fertility, which facilitated the understanding of high fecundity of this invasive insect.

**CONCLUSION:** We reported firstly the Y chromosome of *Bactrocera dorsalis*. Our results will also provide target genes for CRISPR/Cas9 based SIT, leading to the development of novel control strategies against tephritid flies.

## 1 Introduction

Tephritidae is a dipteran family comprising more than 5000 species^1^, dozens of which are invasive and extremely harmful to fruit and vegetable industries^2, 3^. The oriental fruit fly, *Bactrocera dorsalis* (Hendel) attacks more than 600 fruit and vegetable crops^4^, which causes great economic losses^5^. Meanwhile, it is classified as the top member in the competitive hierarchy of tephritid flies^6^, since it could replace and drive various fruit fly species to extinction^7^. The female flies lay eggs inside of the fruits and larvae feed on the flesh, which makes them hard to control^8^. Current management of this fly depends on luring and killing by insecticides^9^, while very much hope has been given to the sterile insect technique (SIT)^10, 11^. The central principle of SIT is to produce a large number of sterile males^12-14^. Male tephritid flies have a unique Y chromosome based on the reported karyotype as 2n = 12 (5A × 2 + X + Y)^15^. According to previous studies, several genes on the Y chromosome are crucial for insect sex determination and male fertility^16-19^, which makes them the most suitable molecular targets for the application of transgenic SIT^20-22^.

Although the genome of female *B. dorsalis* has been reported by several groups^23-25^ including our own group^26^, the Y chromosome of *B. dorsalis* is not available yet. There are several reasons that the Y chromosomes of insects are extremely difficult to assemble. Most highly degenerated Y chromosome has experienced several rounds of recombination suppression^27-29^, leading to repeat accumulation and heterochromatinization^30-34^. Earlier attempts in analyzing of insect Y chromosomes have shown an extremely large proportion of satellite DNA and transposons elements^32, 35-38^. Sex chromosome differentiation also leads the Y chromosome to be hemizygous, which reduces its sequencing coverage^39, 40^. The lack of Y chromosome assembly prevents in-depth understanding the contribution of Y chromosome to the sex determination and male fertility, as well as the development of Y chromosome based transgenic SIT strategy to control this notorious fly.

In this study, we assembled the Y chromosome sequence of *B. dorsalis*, which is the first reported Y chromosome in Tephritidae. The assembled Y chromosome contains 61 genes. We surprisingly found that the M factor (*MoY*), widely conserved in Tephritidae^41^,is a multi-copy gene. In addition, *gyf-like on the Y chromosome* (*gyfY*) has experienced dramatic gene amplification in several *Bactrocera* species. By RNAi mediated gene knockdown, we found that the amplification of *gyfY* is essential for male fertility in *B. dorsalis*. This facilitated the understanding of high fecundity of this invasive insect. Our results will also provide target genes for CRISPR/Cas9 based SIT, which may lead to the development of novel control strategies against the tephritid flies.

## 2 Materials and methods

### 2.1 Flies

The *Bactrocera dorsalis* population sequenced in the present study was maintained since a wild population sampled in Haikou, Hainan province, China, in 2008. Adults were reared under standard conditions (27 ± 1 °C, 70 % ± 5 % relative humidity, and a 14-h photoperiod), on an artificial diet (containing honey, sugar, yeast powder, vitamin C, and water). A male fly from the inbred line at the 7^th^ generation was collected for genomic DNA extraction. To avoid contamination by food and the intestinal microorganism, proboscis and the entire abdomens of the adults were removed and the surfaces were treated with 75 % ethanol.

### 2.2 Genome sequencing

Genomic DNA used for Pacbio sequencing was extracted via the sodium dodecyl sulfate (SDS) method. We used the PacBio Sequel sequencing platform by Beijing Novogene Bioinformatics Technology Co., Ltd. (Novogene, Beijing, China) platform to achieve deep coverage of sequencing reads. Libraries were constructed by SMRTbell^®^ Express Template Preparation Kit 2.0 (Pacific Biosciences, Delaware, USA) according to the manufacturer’s instructions^42^. For short-read sequencing, an Illumina library with an average insert size of 350 bp was constructed using NEB Next Ultra DNA Library Prep Kit (NEB, Beijing, China) with PE 150 bp mode.

Hi-C sequencing data was used to improve draft genome assemblies and assist the chromosome-level assembly^43^. DNA from 25 male flies without the entire abdomens and proboscis was extracted and randomly broken into segments with 350 bp for Hi-C library preparation. Hi-C sequencing was conducted on the Illumina Novaseq 6000 platform with the PE 150 bp mode.

DNA used for Bisulfite-seq was extracted from eggs of *B. dorsalis* via Magnetic Universal Genomic DNA Kit (TIANGEN, Beijing, China) following the manufacturer’s recommendations. After bisulfite treatment, sequencing Illumina library construction of the size selected samples was performed by Accel-NGS^®^ Methyl-Seq DNA Library Kit (Swift, MI, USA) with PE 150 bp mode. Bismark (v0.22.1)^44^ was used for mapping, deduplication and methylation extraction.

### 2.3 Chromosome-level assembly and genome assessment

Firstly, to estimate genome size and heterozygosity, a *k*-mer analysis (jellyfish v2.2.10)^45^ was carried out by clean HiFi data and visualized using GenomeScope2 (v2.0)^46^. Hifiasm (v0.17.7)^47^ with Hi-C integration mode was used for genome assembly. The heterozygosity of the assembly was removed by Purge_dups (v1.2.5) with default parameters^48^. The Hi-C sequencing reads were then mapped to the contigs using the Juicer (v1.6) pipeline^49^. Following this, the 3D-DNA (v201013) pipeline^50^ was used to generate the “.hic” file. Subsequently, we visualized the Hi-C heatmap using Juicebox Assembly Tools^51^ for manual operation. According to the Hi-C interaction heatmap in Juicebox. We demarcated chromosome boundaries and adjusted the contig order to acquire a high-quality chromosome-level genome assembly (Figure S1). Finally, Benchmarking Universal Single-Copy Orthologues (BUSCO v5.5.0)^52^ with the insecta_odb10 lineage (n = 1367) was used to assess genome assembly completeness.

### 2.4 Identifying the sex chromosomes and autosomes with re-sequencing data

The chromosome quotient method^53^ was used to identify the sex chromosomes. In detail, the female and male re-sequencing reads were mapped to the assembled genome using BWA-mem (v0.7.17-r1188)^54^, and then samtools (v 1.10)^55^ and bedtools (v2.25.0)^56^ is used to calculate the sequencing coverage and depth in 100k windows. The ratio of female to male depth values is expected to cluster around 0, 1 or 2 for Y, autosomal or X chromosome respectively.

### 2.5 Repeat and genome annotation

Insect homology repetitive elements were obtained from RepeatMasker (http://www.repeatmasker.org) database (RepeatMaskerLib.h5) and Repbase (RepeatmaskerEdition-20181026)^57^. Satellite Repeat Finder (SRF)^58^, RepeatModeler (v2.0.3) and EDTA (v2.2.0)^59^ were used for de novo prediction of repetitive elements. RepeatMasker (v4.1.5) was utilized to annotate the repeat elements. For tandem repeats, TideHunter (1.4.2)^60^ was used to predict the repeat units and StainedGlass (v0.1)^61^ was used for visualization.

We subjected Hisat2 (v2.2.1)^62^ to align RNA-seq data from 11 tissues (Table S3) and Trinity (v2.8.5)^63^ to assemble transcripts. Maker (v3.01.03)^64^ was used to predicted gene models based on assembled transcripts and protein homology based on the *Zeugodacus cucurbitae, Drosophila melanogaster*^65^ and female *B. dorsalis*^24, 26^ generated by liftoff (v1.6.3)^66^. Finally, we used PASA (v2.5.2)^67^ pipeline to polish gene models using Trinity assembly.

For genes on the Y chromosome, we manually analyzed and removed some fake genes meeting the following four criteria that 1, the cds length of the putative gene is shorter than 250 bp; 2, the FPKM value is almost zero; 3, the number of exons not more than two; 4, lacking orthologous gene with the proximal species.

### 2.6 Determination of *gyfY* and *MoY* copy number using ddPCR analysis

Genomic DNA was extracted from the individual fly using TIANamp Genomic DNA kit (TIANGEN, Beijing, China) and was digested by the restriction enzymes MboI. Copy number of Y-link genes *gyfY* and *MoY* were assessed via ddPCR using a QX200 Droplet Digital™ PCR System (Bio-Rad, CA, USA) in accordance with the manufacturer’s protocol of ddPCR Supermix for Probes (No dUTP) (Bio-Rad, CA, USA). Reference gene *IR25a*^68^ was fluorescently labelled with VIC, while target genes were labelled with FAM (Table S7). In brief, each 22 μL reaction contained 11 μL of 2 × ddPCR Supermix, 0.99 μL each of 10 μM primers, 0.55 μL each of 10 μM probes and 0.55 ng genomic DNA. About 40 μL of droplet reaction was generated in the QX200 Droplet Generator with the assistance of droplet generation oil. The PCR were performed following cycling conditions: 1 × (95 °C for 10 min), 40 × (94 °C for 30 s, 57 °C for 1 min), 1 × (4 °C for 30 min). Following amplification, PCR plate 96 (Eppendorf, Hamburg, Germany) was transferred to the QX200 Droplet Reader and analyzed using QuantaSoft to acquire the absolute quantification (Table S8, Figure S5).

### 2.7 RNAi of *gyfY* and male sperm viability

A long dsRNA (1056 bp) covering all the copies of the *gyfY2* was designed to knockdown the expressions of all *gyfY2* copies, while a specific dsRNA was designed targeting the *gyf on the X chromosome* (*gyfX*) (The primers are listed in Table S7). The dsRNAs were synthesized with the TranscriptAid T7 High Yield Transcription Kit (Thermo Scientific, DE, USA). Two micrograms of dsRNA (ds*GFP* was used as a negative control) was injected to the immature male twice (in 1-day-old and 5-day-old, respectively) to obtain better silence efficiency with a standard protocol^69^. Then, the mature males were dissected and the testes were collected for the viability assay in 9-day-old.

The spermatozoa were diluted in a buffer containing bovine serum albumin (10 mM HEPES, 2 % BSA, 150 mM NaCl, pH 7.4), and three pairs of testes were dissected in 100 mL buffer. According to the manufacturer’s instructions, the LIVE/DEAD™ Sperm Viability Kit (Thermo Scientific, MA, USA) was used to dye sperms. In detail, two luminous dyes, SYBR-14 and PI, were applied to distinguish live and dead sperms, respectively^70^. The number of sperms was counted by photographing under the stereomicroscope (Leica, Wetzlar, Germany). Following the process above, the specific ds*gyfX* was used for RNA interference *gyfX*.

## 3 Results

### 3.1 A reference genome assembly of a male *Bactrocera dorsalis*

About 19.3 GB PacBio HiFi data was obtained from an individual male fly. With the assistance of about 45.4 GB Hi-C data (Table S2), we generated a raw assembly comprising 631 MB. Purge_dups (v1.2.5)^48^ was used to remove the 67.9 Mb redundant contigs. Compared with the *k*-mer based genome size estimation (495 Mb) (Figure 1a), we uncovered 67.6 Mb more sequences, in part due to succesful assembly of some highly repetitive satellite DNA. The final BUSCO completeness score is 98.3 %, with 97.4 % single, 0.9 % duplicated, 0.3 % fragmented, and 1.4 % missing (Figure S2), suggesting the genome is highly completed.

**Figure 1.**
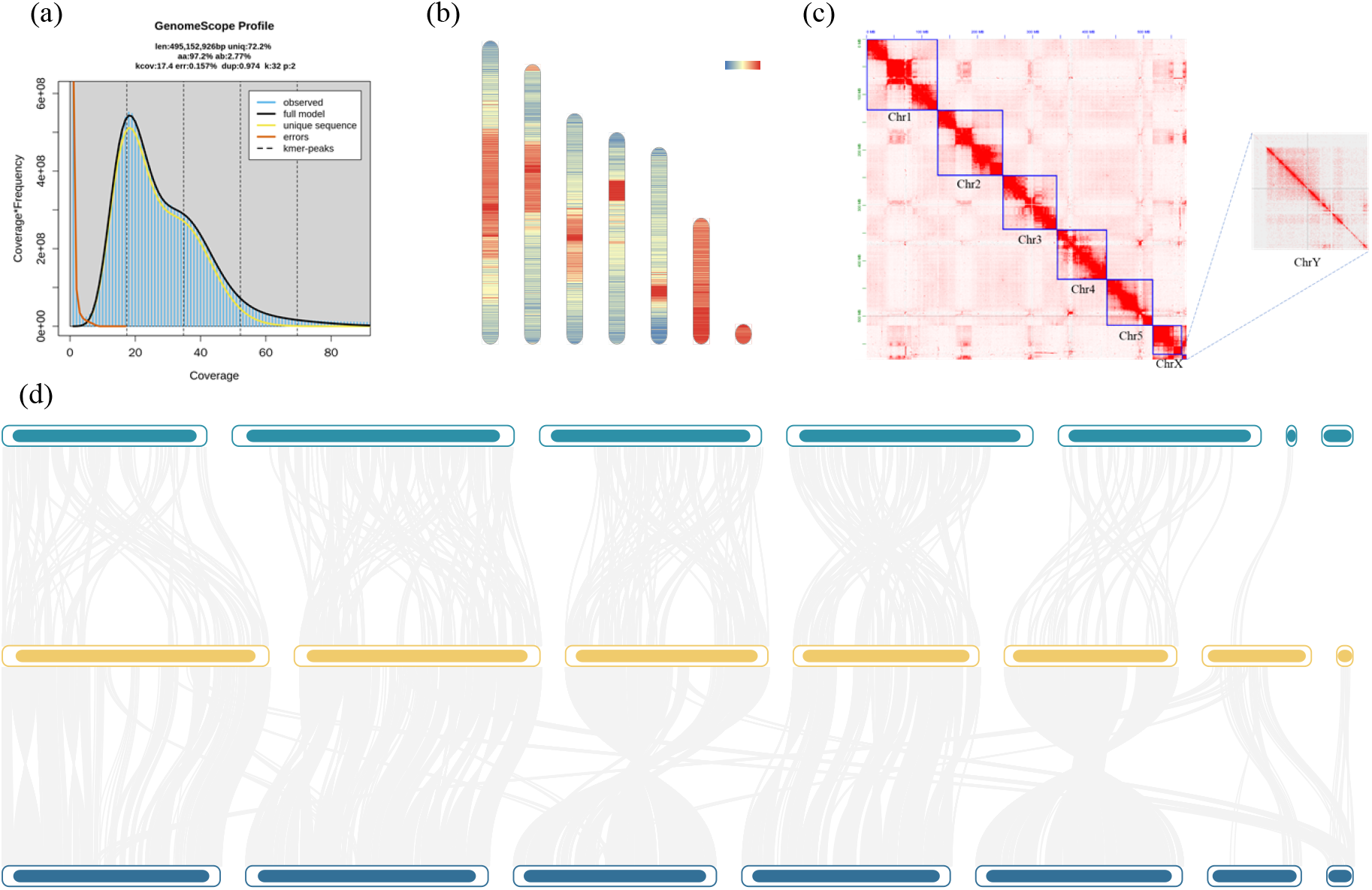
Chromosome-level genome assembly features of *Bactrocera dorsalis*. (a) Estimation of the genome size based on the distribution of *k*-mers (*k* = 32) of HiFi reads. (b) Chromosome-level genome repeat sequence characteristics. (c) Hi-C heatmap for the whole genome and zoom-in view for Y chromosomes visualised in Juicebox. (d) Whole-genome synteny between *Z. cucurbitae* (Zcuc), *D. melanogaster* (Dmel) and *B. dorsalis* (Bdor).

We further anchored the contigs to chromosome level with the assistance of Hi-C data, generating the final assembly comprising 562.6 MB, with a scaffold N50 of 93.2 MB and contig N50 of 2.6 MB (Table S4). The Hi-C heatmap revealed 7 chromosome models (Figure 1c), including 5 autosomes, X chromosome and Y chromosome, accounting for 98.1 % of the assembled sequence. Compared with other published Tephritidae genomes (Table S1), the genome of *B. dorsalis* is the most completed and the best assembled with the highest scaffold N50.

Our high-quality assembly contains large amounts of highly repetitive regions, including pericentromeric regions, putative centromeric repeats and the repeat-rich sex chromosomes (X and Y chromosomes). Overall, about 310 Mb of the assembled 563 Mb *B. dorsalis* genome are repetitive (Table S6). It is noted that SINEs (short interspersed elements) are almost absent from of *B. dorsalis*, despite the overall high repeat content. After masking the repeat sequences, a total of 16,773 protein-coding genes were annotated.

### 3.2 Centromere

Our Hi-C interaction map shows strong inter-chromosomal interactions among centromeres and pericentromeric regions (Figure 1c), similar to that in *D. melanogaster*^71^. The centromeres and pericentric regions also exhibit the highest repeat content and methylation and lowest gene density across the chromosomes (Figure 1b, S4). The pericentric regions appear to be larger on larger chromosomes. For instance, in Chr1, the pericentric regions occupies almost one-third of the entire chromosome (Figure 1c). Our syntenic analysis shows that those pericentric regions are not present in *D. melanogaster*, but are partially present in *Z. cucurbitae* (Figure 1d), suggesting graduate expansion of the pericentric repeats. Satellite DNA is the major repeat category in those regions, and is dominated by five different satellite DNA families. Those five families exclusively distribute in the centromeric regions, and are present in all chromosomes (Figure S3).

### 3.3 Characterizations of the Y chromosome

We carried out genome resequencing of male and female flies, producing clean data of about 128.2 Gb and 56.6 Gb, respectively (Table S3). Using these data, we identified X and Y chromosomes in *B. dorsalis* with the chromosome quotient method^53^. The second smallest chromosome (Chr6) has a nearly two-fold greater sequencing depth compared to females, consistent with the expected for the X chromosome. The smallest chromosome (Chr7) has a male-biased coverage and depth ratio and was consistent with the expectation for the Y chromosome (Figure 2a, d). The other 5 chromosomes exhibit no significant differences between females and males, with female/male ratios close to 1 (Figure 2a). According to the repeat annotation, repeat content of the total sequences is various among autosomes, X and Y chromosome, which is 47.86 %, 80.62 % and 74.47 %, respectively (Table S5).

**Figure 2.**
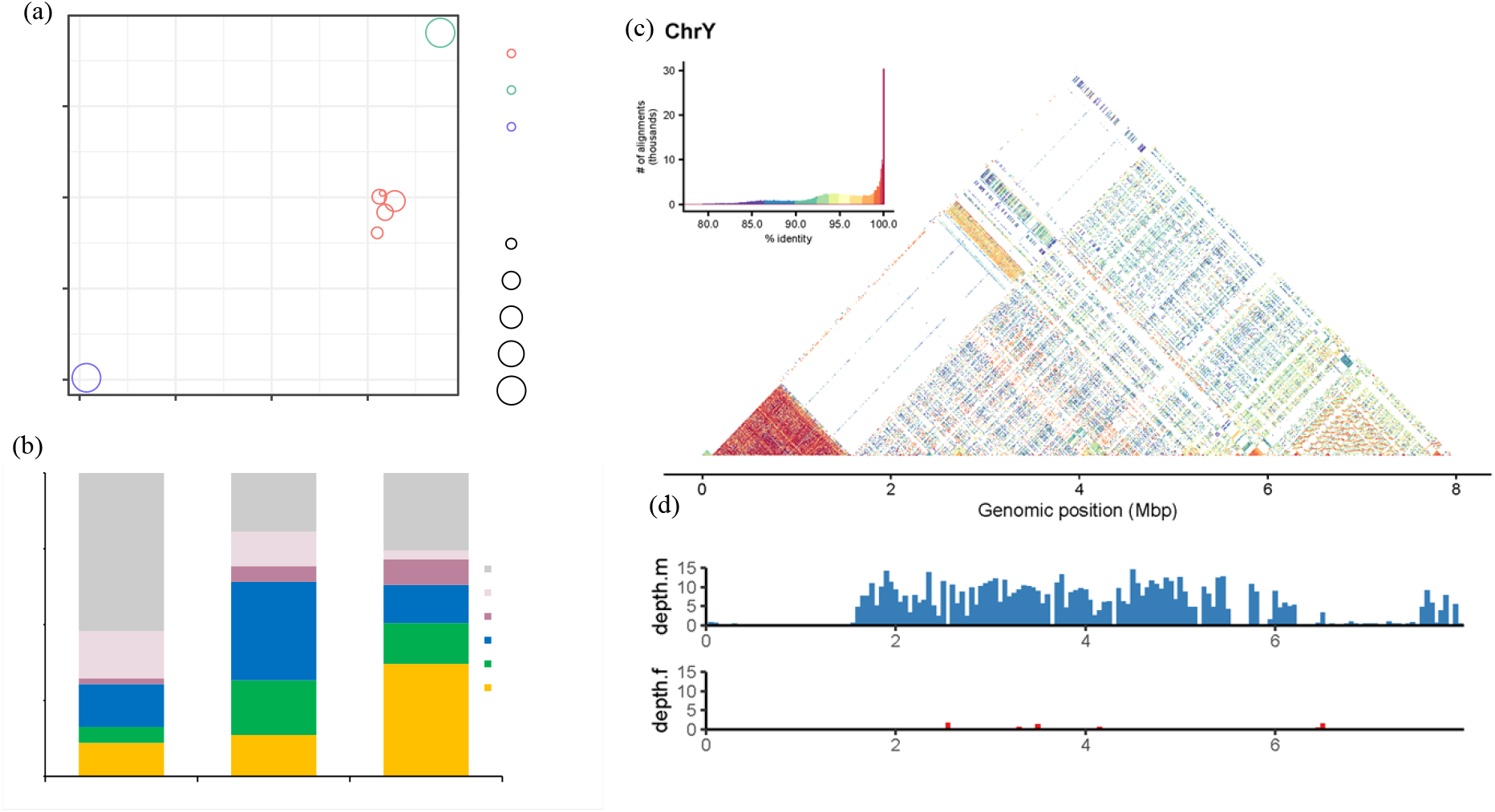
Identification and repeat sequence characteristics of Y chromosome. (a) Female to male depth ratio and coverage ratio for Y chromosome are both close to zero, for X chromosome are close to two and one, respectively. (b) Color-coded bar plot illustrating the proportions of TE compositions in the autosomal, X, and Y chromosomal sequences. (c) Identification of satellite DNA on the Y chromosome by StainedGlass. (d) The bar charts show male (depth.m) and female (depth.f) re-sequencing coverage and depth for Y chromosome.

Most of these repeat sequences in Y chromosome were LINEs (37.07 %), while LTRs and other DNA elements accounted for 13.46 % and 12.56 %, respectively. Compared with autosomes, the rates of LTR and LINEs of Y chromosome are close to 3 (Figure 2b). And the satellite DNA of the Y chromosome is mainly concentrated at the beginning (Figure 2c).

Due to the repetitive nature of the Y chromosome, some bioinformatically predicted gene models may be incomplete or erroneous. To increase the quality of gene annotations, we manually curated gene models, leading to the identification of 61 Y-linked genes (Figure 3). Those Y-linked genes include 9 multi-copy genes, among which 5 were tandemly amplified on the Y chromosome. Besides, we uncovered that the M factor (*MoY*) in Tephritidae has two copies due to tandem duplication, verified by our droplet digital PCR (ddPCP) analysis (Table S8).

**Figure 3.**
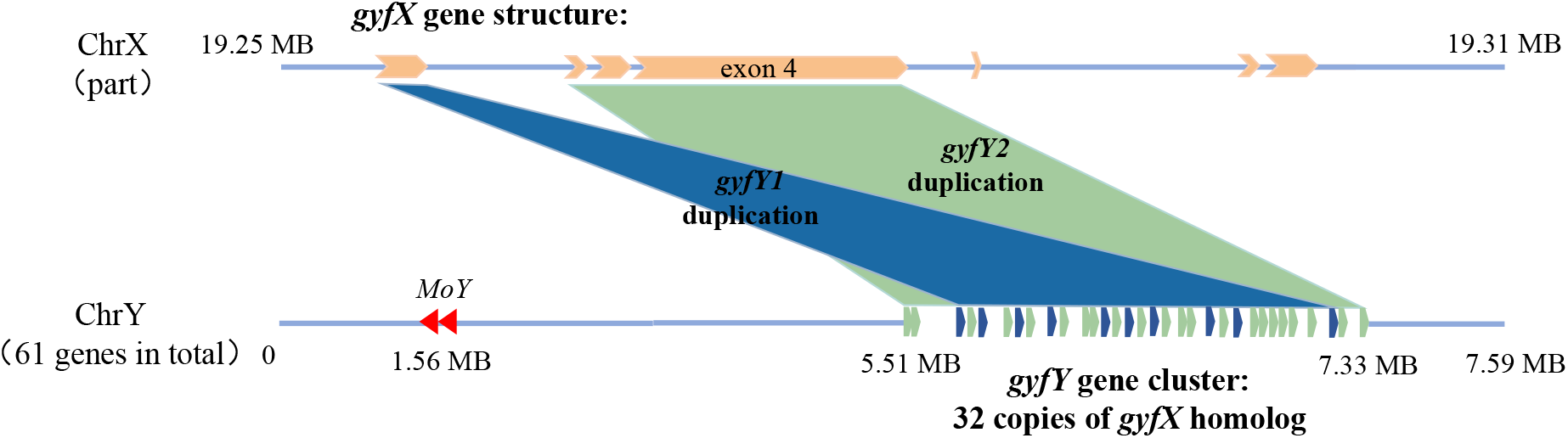
Schematic diagram of *gyfY* and *MoY* gene distribution on the Y chromosome. *MoY* (Red) has two adjacent copies. *gyfY1* (Blue) is homologous to exon 1 of *gyfX*, and *gyfY2* (Green) is homologous to exon 2, 3 and part of 4 of *gyfX*. The *gyfY* genes are all on the positive strand of the Y chromosome. For more details, please see Figure S6, the multiple sequence alignments of the putative gyfY and gyfX proteins.

### 3.4 Male sperm viability after *gyfY* RNAi

gyf (glycine-tyrosine-phenylalanine) protein is ubiquitous in eukaryotes^72, 73^. We found that *gyfY* was amplificated dramatically on the Y chromosome and was identified as a truncated paralog of *gyf on the X chromosome* (*gyfX*) in *B. dorsalis* (Figure 3). The copy number of *gyfY* is 27 (9 copies of *gyfY1* and 18 copies of *gyfY2*) confirmed by ddPCR, that was similar with the prediction by genome annotation (32 copies) (Figure 3, Table S8).

In order to explore the function of the *gyfY*, we designed a dsRNA with a length of 1056 bp, theoretically capable of broadly interfering with all *gyfY2* genes. Male flies were injected twice in 1-day-old and 5-day-old, and then 30 male flies were detected in 9-day-old. We found that the treated males had almost no live sperm compare to controls that had about 20 mature sperms per square millimeter (Figure 4b, d), indicating that the amplification of *gyfY* is essential for male fertility. In addition, RNA interference with *gyfX* also resulted in an about 30% reduction in the number of live sperm (Figure 4c, d).

**Figure 4.**
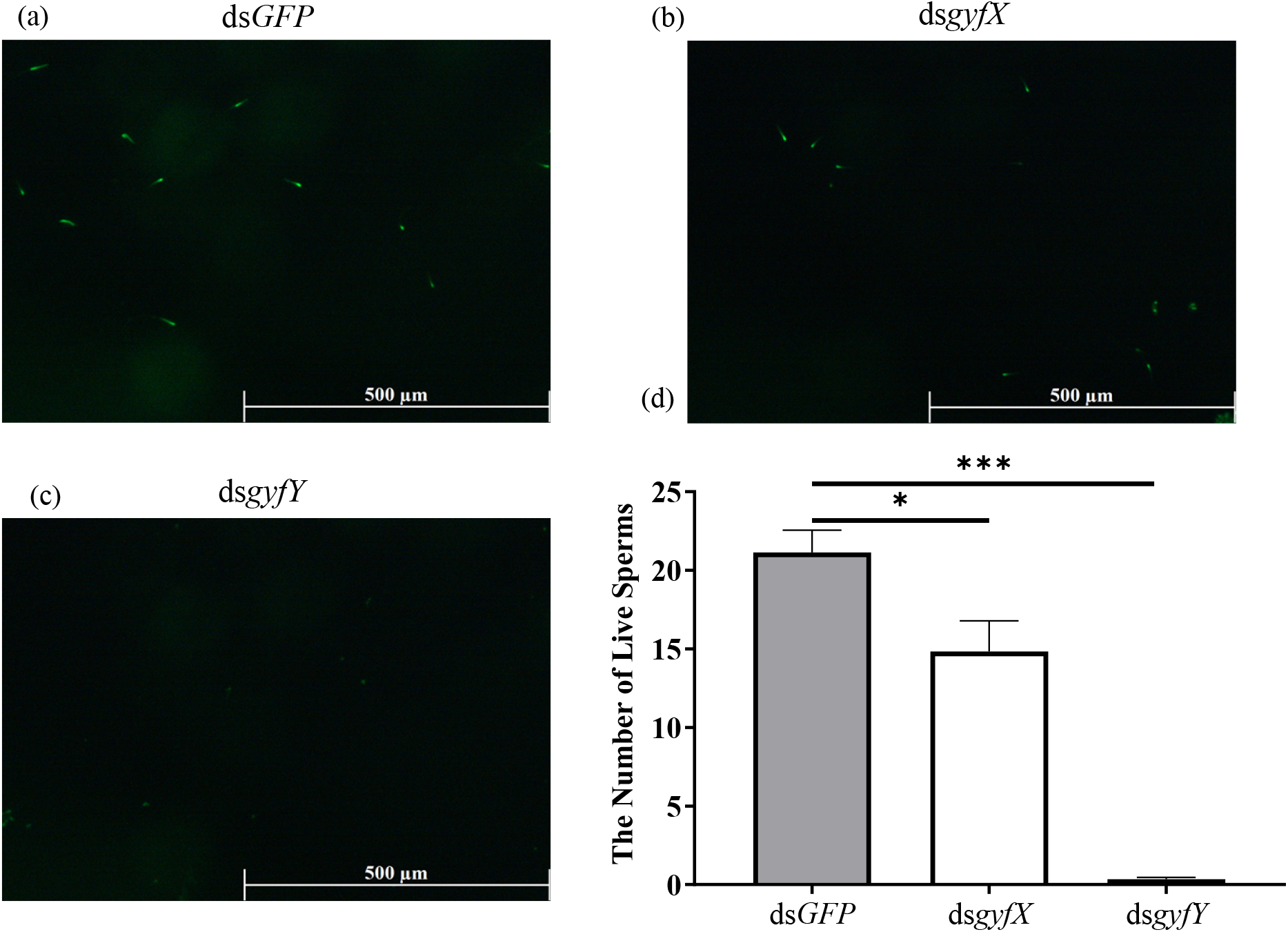
Effect of *gyfX* and *gyfY* RNA interference (RNAi) on sperm viability. (a-c) Live sperms after injection with ds*GFP*, ds*gyfX* and ds*gyfY*. Green indicates live sperms, and scale bars are indicated in the bottom left corner. (d) Number of live sperms after injected twice ds*GFP*, ds*gyfX* and ds*gyfY*. Data were presented as mean ± SE (*n* = 50). Asterisks represent a significant difference determined by Student’s *t*-test (\**P* < 0.05; \*\*\**P* < 0.001)

## 4 Discussion

Male tephritid flies own a unique Y chromosome, which is crucial for insect sex determination and male fertility^18, 41, 74^. However, the Y chromosomes is the most difficult part of the genome, because it is highly heterochromatic and repetitive^75, 76^, and has low sequence coverage^40^. Using the PacBio HiFi, Hi-C and re-sequencing data, we generated the chromosome-level genome assembly of male *B. dorsalis*, including a characterized Y chromosome. This coordinated Y chromosome of *B. dorsalis* is the first assembled Y chromosome reported in tephritid flies. The completion of the Y chromosome assembly in tephritid flies is significant for understanding the intricacies of genome function and evolution.

The overall TE content of tephritid flies range from 21.87 % to 55.03 %, according to the available genomic information (Table S1, Table S9). This likely contributes to the genome size variation of the flies. Based on our current assembly, we found that the overall TE content of *B. dorsalis* (55.03 %) is much higher than *D. melanogaster* (21.53 %)^65^. However, no SINE family has been detected in *B. dorsalis*, which also occurs in most insects^77^. The repeat sequence content of X and Y chromosomes is 80.62 % and 74.47 % respectively, which is significantly higher than that of autosomes (Figure 2b, Table S5). While the expansion of repeats on the Y chromosome is consistent with prediction due to the loss of recombination^38, 78^, it remains a puzzle about the repeat expansion on the X chromosome. It is possible that the X chromosome has a reduced selection efficacy to purge repeats due to its smaller effective population size compared with autosomes^79^.

The high-quality genome assembly allowed us to identify the centromere and the repetitive pericentric regions of *B. dorsalis* which we found were much larger than those of *Drosophila*^76, 80, 81^. The cause and consequence of pericentric expansion in *B. dorsalis* warrants future epigenetic investigations. Moreover, we identified the putative centromere tandem duplication sequences, which primarily consist of 5 sequences ranging from 165 bp to 180 bp (Figure S3), but to determine the core centromeric repeat, CENP-A ChIP-seq data is required. To the best of our knowledge, this is the first exploration of centromeric sequences in an agricultural insect. Our study provides a working protocol to identify centromeric repeats in insects, which will be useful to illuminate the origin and evolution of centromeric sequences.

Interestingly, we first confirmed the *M factor* (*MoY*) in Tephritidae are multi-copy gene by ddPCP (Table S8). This is in accordance with the prediction of 5 copies of *MoY* on the Y chromosome in *Z. cucurbitae*. However, there is only one copy of M factor in other Dipteran insects, such as mosquitos *Anopheles stephensi, Aedes aegypti* and *Anopheles gambiae* and housefly *Musca domestica* (Table S9)^41, 82-85^. To masculinize *B. dorsalis* and the *Z. cucurbitae*, multiple M factors are required to work together or each M factor can work separately. Further study on the function of different copies of the *MoY* will provide a clue why multiple copy number of *MoY* is needed in the tephritid flies.

gyf (glycine-tyrosine-phenylalanine) proteins are characterized by the presence of a conserved domain (Pfam domain PF02213), which is ubiquitous in eukaryotes^72, 73^. In *D. melanogaster, gyf* is on the dot chromosome (homologous to the X chromosome of Tephritidae)^86^, which functions in regulation of autophagy and mRNA regulatory element binding as a translation repressor^87, 88^. In the current study, a copy of *gyf* (*gyfX*) was predicted on the X chromosome, while 32 copies of *gyf-like on the Y chromosome* (*gyfY*) were predicted on the Y chromosome of *B. dorsalis* (Figure 3). This dramatical amplification of *gyfY* may be caused by non-allelic homologous recombination to enhance the expression^89^. The functional domains of *gyfX* were intact and well conserved comparing to other eukaryotes species. However, multicopy *gyfY* is identified as the truncated paralog of *gyfX*.

We identified two types of *gyfY* tandem repeats namely *gyfY1* (derived from the first exon of *gyfX*) and *gyfY2* (derived from the second, third and part of fourth exons of *gyfX*), instead of only *gyfY2* in a previous study^18^. Furthermore, we confirmed 9 out of 10 predicted copies of *gyfY1*, while confirmed 18 out of the 22 predicted copies of *gyfY2*. This is in accordance with the previous report, while the few unconfirmed copies may be due to the complexity and repetition of the Y chromosome. Based on the analysis of the genome information available, similar gene amplification of *gyfY* occurs on the Y chromosome of *Bactrocera tryoni*^90^, but surprisingly not in *Bactrocera oleae*^91^ or *Z. cucurbitae*. This might be due to the rapid differential evolution of the Y chromosome^38^. It is noticeable that *gyfY* has the domain of disorder protein but no PF02213 domain which was also verified by the PONDR (Predictor of Natural Disordered Regions; http://pondr.com/) (Figure S7)^92^.

A long ds*gyfY* (1056 bp) was designed from the region of the *gyfX* that is homologous to the *gyfY2*. The ds*gyfY* was decomposed into many siRNAs to knockdown all the *gyfY2*. With the 73% silence efficiency (Figure S8), the flies treated with ds*gyfY* almost had no live sperm (Figure 4), which indicated that the explosive amplification of *gyfY* enhanced the male fitness in *B. dorsalis*. Since the long dsRNA covers all the copies of *gyfY2*, our RNAi showed more significant reduction on the number of the live sperms in the male flies than a former study, that live sperms decreased by ∼ 50% due to knocking down a *gyfY* with a 237 bp dsRNA^18^. Moreover, similar gene amplification on Y chromosome occurs and is crucial for male fertility in *D. melanogaster*. Y-link *Mst77Y* undergoes amplification from the autosomal gene *Mst77F* to the Y chromosome, collectively participating in the process of regulating sperm nuclear compression^93^. In addition, when knocking down the *gyfX* resulted in an about 30% reduction of the live sperms (Figure 4). This suggested that *gyfX* also have an effect on sperm viability in *B. dorsalis* but less significant than *gyfY*.

In conclusion, we successfully assembled the Y chromosome of *B. dorsalis*, which is the first reported Y chromosome in Tephritidae. The assembled Y chromosome contains 61 genes, including 9 multi-copy genes. We surprisingly found that the M factor (*MoY*), widely conserved in Tephritidae, is a multi-copy gene. In addition, amplification of *gyf* occurred and plays important roles in the male fertility in this species. Our data revealed male specific genes involved in male fertility which are the ideal molecular targets to develop novel transgenic control methods against this notorious fly, such as the genetic sexing strains targeting Y chromosome. Furthermore, CRISPR/Cas9 targeting Y-link multi-copy *gyfY* may lead to “Y-shredding”, resulting in female-only offspring.

## Supporting information

Supplemental Table S1 to S9, and will be used for the link to the file on the preprint site.

Supplemental Figure S1 to S8, and will be used for the link to the file on the preprint site.

## ACKNOWLEDGEMENTS

We thank Dr. Yang Yang and Mr. Changhao Liang from Southwest University and Miss Xing Li from Yangzhou University for assistance on bioinformatics analysis. We also thank Prof. Marc F. Schetelig and Prof. Ying Yan from Justus-Liebig-University Giessen for their comments on an earlier version of this manuscript. This research was supported by funding from the National Key R&D Program of China (2022YFC2601000), National Natural Science Foundation of China (U21A20222, 32072491, 32370445), 111 Project (B18044) and the China Agriculture Research System of MOF and MARA.

## AUTHOR CONTRIBUTIONS

Hongbo Jiang and Luohao Xu designed research; Shuangxiong Wu, Jiahong Wu, Xinrui Jiang and Luohao Xu worked on analysis of the genome assembly and annotation; Shuangxiong Wu, Quan Lei, Donghai He, Chao Ye, and Dong Wei worked on the functional verification of gyfY; Shuangxiong Wu and Hongbo Jiang wrote the paper; Luohao Xu and Hongbo Jiang revised the manuscript.

## CONFLICT OF INTERESTS

The authors declare no conflict of interests.

## DATA AVAILABILITY

All raw data from this study has been deposited into the China National Center for Bioinformation (CNCB, https://www.cncb.ac.cn/) with the BioProject accession PRJCA026299.

---

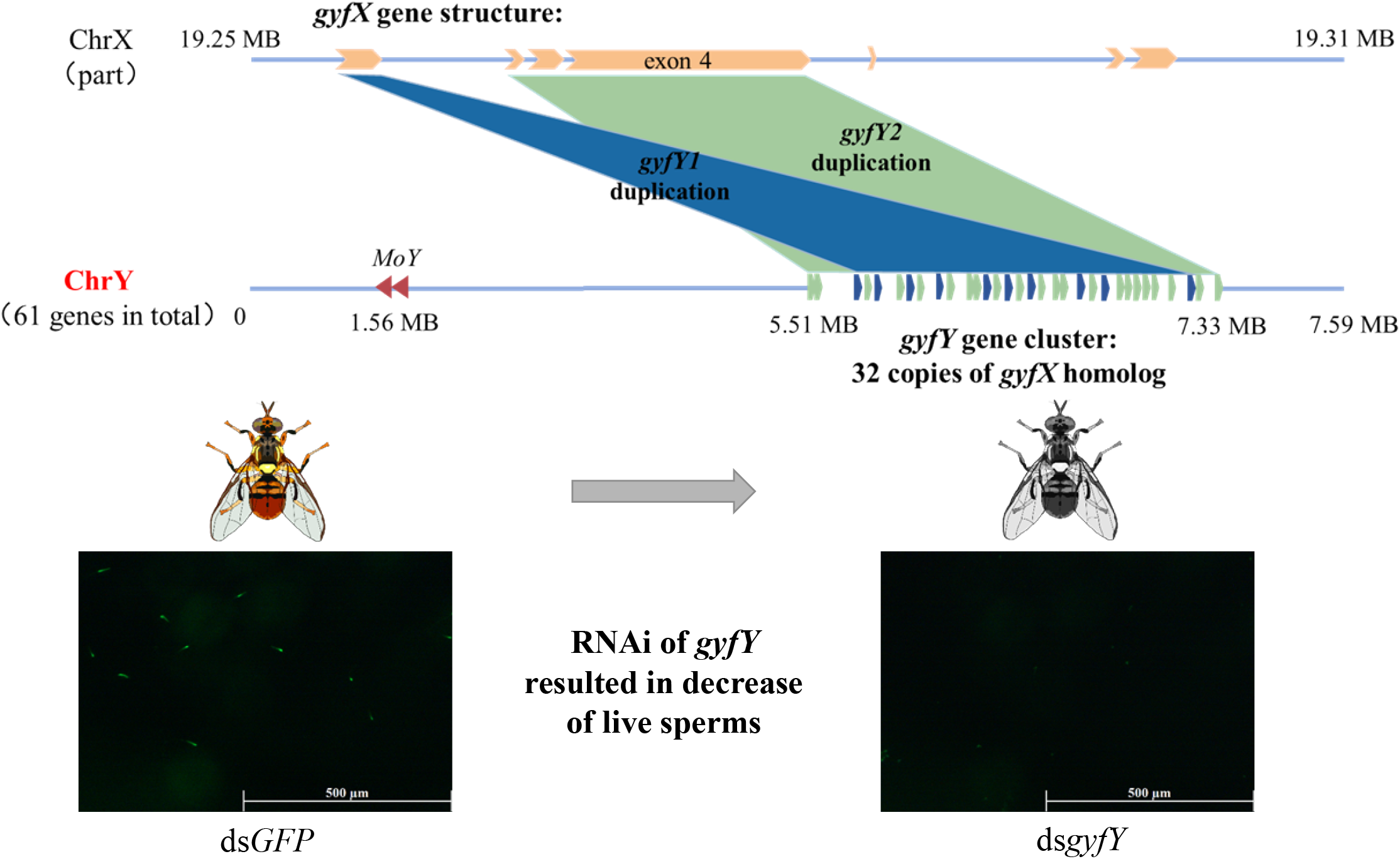

## Notes

### Competing Interest Statement

The authors have declared no competing interest.

