## Supplemental Table S1 to S9, and will be used for the link to the file on the preprint site. for "The assembly of Y chromosome reveals amplification of genes regulating male fertility in *Bactrocera dorsalis*"

**Supplementary Tables**

**Tables S1.** Summary of the genome assemblies of tephritid flies

| | **Bdor_M** | **Bdor_F** | **Btry** | **Bole** | **Bmin** | **Zcuc** | **Ccap** |
|---|---|---|---|---|---|---|---|
| Genome size (Mb) | 562.6 | 524 | 570.6 | 484.9 | 323.8 | 439.2 | 436.5 |
| Gender | male | female | male | male | F+M | male | - |
| Number of contigs | 667 | 702 | 8,397 | 48,617 | 249 | 58 | 3,242 |
| Number of scaffolds | 44 | 370 | 5,891 | 38,160 | - | 55 | 2,354 |
| Number of assembled chromosomes | 5A+X+Y | 5A+X | 5A | - | - | 5A+X+Y | - |
| **Genome assembly quality** | | | | | | | |
| Contig N50 (Mb) | 2.6 | 12.2 | 0.351 | 0.188 | 2.7 | 74.8 | 0.846 |
| Scaffold N50 (Mb) | 93.2 | 90.2 | 81.9 | 4.6 | - | 75.5 | 1.7 |
| **Genomic features** | | | | | | | |
| Repeat percent (%) | 55.03 | 53.05 | 51.55 | 39.09 | 21.87 | 42.97 | 50.54 |
| GC percent (%) | 36.4 | 36.4 | 36.5 | 34.5 | 35 | 35.5 | 35 |
| **Gene annotation** | | | | | | | |
| Number of genes | 16773 | 15379 | 15659 | 15622 | - | 16560 | 19250 |

[#]Bdor_M: male *Bactrocera dorsalis*; Bdor_F: female *Bactrocera dorsalis*; Btry: *Bactrocera tryoni*; Bole: *Bactrocera oleae*; Bmin: *Bactrocera minax*; Zcuc: *Zeugodacus cucurbitae*; Ccap: *Ceratitis capitata*

**Tables S2.** Statistics of sequencing data for *Bactrocera dorsalis* genome assembly

| Sequencing item | Type | Total clean data | Coverage (X) |
|---|---|---|---|
| PacBio HiFi | Subreads | 18.0 Gb | 32.7 |
| Hi-C | PE | 42.3 Gb | 77.0 |

**Tables S3.** Statistics of genomic resequencing and bisulfite sequencing data and transcriptome sequencing of 11 tissues.

| Sample | BaseSum | Q30[#1] | Accession[#2] | Reference |
|---|---|---|---|---|
| **Genomic resequencing** | | | | |
| Female1 | 28.3 Gb | 91.9 % | CNP0003192 | (Yang et al. 2022) |
| Male1 | 19.5 Gb | 93.5 % | This study | |
| Male2 | 19.1 Gb | 93.8 % | This study | |
| Male3 | 25.5 Gb | 93.2 % | This study | |
| **Bisulfite sequencing** | | | | |
| Eggs | 15.2 Gb | 89.9 % | This study | |
| **Transcriptome sequencing** | | | | |
| Brain | 7.0 Gb | 91.6 % | CNP0004721 | (Yang et al. 2024) |
| Testis | 7.7 Gb | 91.7 % | CNP0004721 | (Yang et al. 2024) |
| Fat body | 7.3 Gb | 92.0 % | CNP0004721 | (Yang et al. 2024) |
| Gut | 7.6 Gb | 92.2 % | CNP0004721 | (Yang et al. 2024) |
| Malpighian tubule | 6.8 Gb | 91.5 % | CNP0004721 | (Yang et al. 2024) |
| Ovary | 7.5 Gb | 91.7 % | CNP0004721 | (Yang et al. 2024) |
| Antenna | 6.7 Gb | 93.6 % | CNP0003334 | (Xu et al. 2023) |
| Head cuticle | 6.6 Gb | 93.8 % | CNP0003334 | (Xu et al. 2023) |
| Legs | 6.6 Gb | 93.8 % | CNP0003334 | (Xu et al. 2023) |
| Maxillary palp | 6.7 Gb | 93.7 % | CNP0003334 | (Xu et al. 2023) |
| Proboscis | 6.0 Gb | 93.7 % | CNP0003334 | (Xu et al. 2023) |
| Ovipositor | 6.7 Gb | 94.5 % | CNP0003334 | (Xu et al. 2023) |
| Wings | 6.5 Gb | 94.2 % | CNP0003334 | (Xu et al. 2023) |

[#1]The percentage of bases with Phred quality score ≥ 30

[#2]China National GeneBank DataBase (CNGBdb) with the Project accession number

**Tables S4.** The statistics of contig-level assemblies.

| Stat Type | Raw contigs | Purged contigs |
|---|---|---|
| **Contig number** | 812 | 667 |
| **Contig L50** | 66 | 58 |
| **Contig N50** | 2,615,956 bp | 2,737,110 bp |
| **Longest contig length** | 16,095,034 bp | 16,095,034 bp |
| **Shortest contig length** | 10,824 bp | 15,069 bp |
| **Total bases** | 661,289,832 bp | 589,617,899 bp |

**Tables S5.** Detail information of *Bactrocera dorsalis* genome assembly result by Hi-C.

| Chr | Scaffold number | Length (bp) | GC | Repeat content |
|---|---|---|---|---|
| Chr1 | 117 | 128,467,906 | 36.8 % | 56.00 % |
| Chr2 | 70 | 118,454,170 | 36.5 % | 46.97 % |
| Chr3 | 59 | 97,676,735 | 36.5 % | 44.01 % |
| Chr4 | 49 | 89,628,578 | 36.1 % | 40.63 % |
| Chr5 | 43 | 83,363,771 | 36.0 % | 38.15 % |
| ChrX | 225 | 53,417,179 | 36.1 % | 81.45 % |
| ChrY | 16 | 7,962,575 | 37.7 % | 79.98 % |

[#]Unplaced scaffolds: 10,948,925 bp (account for 1.85 % of the whole genome)

**Tables S6.** Repeat elements annotation

| Type | Element number | Length (bp) | percentage of genome |
|---|---|---|---|
| **Type I: Retroelements** | 204,029 | 110,950,666 | 18.79 % |
| SINE | 0 | 0 | 0 |
| LINE | 151,303 | 69,009,236 | 11.69 % |
| LTR | 52,726 | 41,941,430 | 7.10 % |
| **Type II: DNA transposons** | 278,825 | 93,608,029 | 15.86 % |
| **Type III: Tandem repeat** | 196,811 | 9,247,097 | 1.56 % |
| Satellites | 1,235 | 64,615 | 0.01 % |
| Simple repeats | 171,106 | 7,854,715 | 1.33 % |
| Low complexity | 24,470 | 1,327,767 | 0.22 % |
| **Type IV: Small RNA** | 1,122 | 2,819,481 | 0.48 % |

[#]Unclassified: 13.69 %

**Tables S7.** Primer sequences used in the present study.

| Primer name | Sequence (5' – 3') |
|---|---|
| **Primers for RNAi** | |
| ds*gyfY*-F | taatacgactcactatagggAGTGAAATGCGAACCCCCAA |
| ds*gyfY*-R | taatacgactcactatagggGCTGCGGTTGCTGCTAATAA |
| ds*gyfX*-F | taatacgactcactatagggTGTTCCACCCTTACCAGTGC |
| ds*gyfX*-R | taatacgactcactatagggTCCGTCATTTCGATCGCTGT |
| ds*GFP*-F | taatacgactcactatagggCAGTTCTTGTTGAATTAGATG |
| ds*GFP*-R | taatacgactcactatagggTTTGGTTTGTCTCCCATGATG |
| **Primers for Quantitative Real-time PCR** | |
| q*gyfY*-1-F | GAAATATTTGTGGAGAAGGTGCACTT |
| q*gyfY*-1-R | AGGTATTTTCTTAATCTAGAGGACCTCCT |
| q*gyfY*-2-F | AGCCGAAGATAAACATCGAGAGAA |
| q*gyfY*-2-R | GTACTAGAATTTGCAGCTGCCGTT |
| q*gyfX*-F | GGATGAGGGAGGTTACTGAC |
| q*gyfX*-R | GCATGGCAGTTACTGTTGTA |
| **Primers and probes for ddPCR** | |
| d*gyfY1*-F | GAAATATTTGTGGAGAAGGTGCACTT |
| d*gyfY1*-R | AGGTATTTTCTTAATCTAGAGGACCTCCT |
| d*gyfY1*-P | AACTGCTGCTACAGGTTCCGTCGGTT |
| d*gyfY2*-F | AGCCGAAGATAAACATCGAGAGAA |
| d*gyfY2*-R | GTACTAGAATTTGCAGCTGCCGTT |
| d*gyfY2*-P | ACGCCACCAACAACAATTAGATGATGAA |
| d*MoY*-F | AAGGAAATAGTACTGAATTTAAGCACA |
| d*MoY*-R | TTCGCAAACTGCATTGATTCTT |
| d*MoY*-P | TTGCTGAATACACAGAAGTCACGGAGCTA |
| d*IR25a*-F | AAGTGGTGGAAAAACGATGAAA |
| d*IR25a*-R | TGTATGGAAATGCCGTCTGACT |
| d*IR25a*-P | ACAAGCCAAATGTGACAAGCCAGAAGAC |

**Tables S8.** Determination of *gyfY* and *MoY* copy number by ddPCR

| Sample | Target | Concentration (copies/µL) | Positive Droplets | Total Droplets | Ratio[#2] |
|---|---|---|---|---|---|
| Female1 | *gyfY1* | 0 | 0 | 14,527 | 0 |
| | *IR25a*[#1] | 27.4 | 313 | | |
| Female2 | *gyfY1* | 0 | 0 | 16,373 | 0 |
| | *IR25a* | 19.2 | 248 | | |
| Male1 | *gyfY1* | 216 | 2,354 | 14,898 | 9.2 : 2[#3] |
| | *IR25a* | 47.2 | 549 | | |
| Male2 | *gyfY1* | 107 | 1,302 | 15,951 | 8.5 : 2 |
| | *IR25a* | 25.2 | 316 | | |
| Male3 | *gyfY1* | 104 | 1,489 | 17,567 | 9.2 : 2 |
| | *IR25a* | 22.6 | 313 | | |
| Female1 | *gyfY2* | 0 | 0 | 15,919 | 0 |
| | *IR25a* | 19.3 | 243 | | |
| Female2 | *gyfY2* | 0 | | 17,212 | 0 |
| | *IR25a* | 20 | 271 | | |
| Male1 | *gyfY2* | 49.9 | 5,649 | 18,882 | 17.9 : 2 |
| | *IR25a* | 447 | 734 | | |
| Male2 | *gyfY2* | 197 | 2,694 | 18,617 | 18.6 : 2 |
| | *IR25a* | 21.2 | 311 | | |
| Male3 | *gyfY2* | 208 | 2,809 | 18,464 | 17.4 : 2 |
| | *IR25a* | 23.9 | 348 | | |
| Female1 | *MoY* | 0 | 0 | 15,010 | 0 |
| | *IR25a* | 32.4 | 382 | | |
| Female2 | *MoY* | 0 | 0 | 17,613 | 0 |
| | *IR25a* | 19.6 | 272 | | |
| Male1 | *MoY* | 44.4 | 567 | 16,337 | 1.9 : 2 |
| | *IR25a* | 46.3 | 590 | | |
| Male2 | *MoY* | 23.3 | 340 | 18,488 | 1.8 : 2 |
| | *IR25a* | 26.1 | 379 | | |
| Male3 | *MoY* | 20.9 | 280 | 16,991 | 1.8 : 2 |
| | *IR25a* | 23.7 | 317 | | |

[#1]*IR25a* is a reference gene.

[#2]Ratio = Concentration of target gene / Concentration of *IR25a*.

[#3]The target genes located on the Y chromosome, and *IR25a* located on the Chr1 with 2 copies

**Tables S9.** Download links for genomic profiles used in this study.

| Species | Abbreviation | URL | Reference |
|---|---|---|---|
| *Bactrocera dorsalis* | Bdor_M | This study | |
| *Bactrocera dorsalis* | Bdor_F | https://ftp.cngb.org/pub/CNSA/data5/CNP0003192/CNS0572595/CNA0050866/scaffold_genome.review6.assembly.FINAL.fasta.gz | (Yang et al. 2022) |
| *Bactrocera tryoni* | Btry | https://ftp.ncbi.nlm.nih.gov/genomes/all/GCA/016/617/805/GCA_016617805.2_CSIRO_BtryS06_freeze2/GCA_016617805.2_CSIRO_BtryS06_freeze2_genomic.fna.gz | (Choo et al. 2019) |
| *Bactrocera oleae* | Bole | https://ftp.ncbi.nlm.nih.gov/genomes/all/GCA/001/188/975/GCA_001188975.4_MU_Boleae_v2/GCA_001188975.4_MU_Boleae_v2_genomic.fna.gz | (Bayega et al. 2020) |
| *Bactrocera minax* | Bmin | https://ftp.ncbi.nlm.nih.gov/genomes/all/GCA/021/498/325/GCA_021498325.1_ASM2149832v1/GCA_021498325.1_ASM2149832v1_genomic.fna.gz | (Wang et al. 2022) |
| *Zeugodacus cucurbitae* | Zcuc | https://ftp.ncbi.nlm.nih.gov/genomes/all/GCA/028/554/725/GCA_028554725.2_idZeuCucr1.2/GCA_028554725.2_idZeuCucr1.2_genomic.fna.gz | - |
| *Ceratitis capitata* | Ccap | https://ftp.ncbi.nlm.nih.gov/genomes/all/GCA/000/347/755/GCA_000347755.4_Ccap_2.1/GCA_000347755.4_Ccap_2.1_genomic.fna.gz | (Papanicolaou et al. 2016) |
| *Drosophila melanogaster* | Dmel | https://ftp.ncbi.nlm.nih.gov/genomes/all/GCA/000/001/215/GCA_000001215.4_Release_6_plus_ISO1_MT/GCA_000001215.4_Release_6_plus_ISO1_MT_genomic.fna.gz | (Hoskins et al. 2015) |
| *Anopheles stephensi* | Aste | https://ftp.ncbi.nlm.nih.gov/genomes/all/GCA/013/141/755/GCA_013141755.1_UCI_ANSTEP_V1.0/GCA_013141755.1_UCI_ANSTEP_V1.0_genomic.fna.gz | (Chakraborty et al. 2021) |
| *Aedes aegypti* | Aaeg | https://ftp.ncbi.nlm.nih.gov/genomes/all/GCA/002/204/515/GCA_002204515.1_AaegL5.0/GCA_002204515.1_AaegL5.0_genomic.fna.gz | (Matthews et al. 2018) |
| *Anopheles gambiae* | Agam | https://ftp.ncbi.nlm.nih.gov/genomes/all/GCA/001/542/645/GCA_001542645.1_ASM154264v1/GCA_001542645.1_ASM154264v1_genomic.fna.gz | - |
| *Musca domestica* | Mdom | https://ftp.ncbi.nlm.nih.gov/genomes/all/GCA/002/191/195/GCA_002191195.1_ASM219119v1/GCA_002191195.1_ASM219119v1_genomic.fna.gz | (Meisel, Gonzales, and Luu 2017) |

Hoskins, Roger A., Joseph W. Carlson, Kenneth H. Wan, Soo Park, Ivonne Mendez, Samuel E. Galle, Benjamin W. Booth, Barret D. Pfeiffer, Reed A. George, Robert Svirskas, Martin Krzywinski, Jacqueline Schein, Maria Carmela Accardo, Elisabetta Damia, Giovanni Messina, María Méndez-Lago, Beatriz de Pablos, Olga V. Demakova, Evgeniya N. Andreyeva, Lidiya V. Boldyreva, Marco Marra, A. Bernardo Carvalho, Patrizio Dimitri, Alfredo Villasante, Igor F. Zhimulev, Gerald M. Rubin, Gary H. Karpen, and Susan E. Celniker. 2015. 'The Release 6 reference sequence of the *Drosophila melanogaster* genome', *Genome Research*, 25: 445-58.

Matthews, Benjamin J., Olga Dudchenko, Sarah B. Kingan, Sergey Koren, Igor Antoshechkin, Jacob E. Crawford, William J. Glassford, Margaret Herre, Seth N. Redmond, Noah H. Rose, Gareth D. Weedall, Yang Wu, Sanjit S. Batra, Carlos A. Brito-Sierra, Steven D. Buckingham, Corey L. Campbell, Saki Chan, Eric Cox, Benjamin R. Evans, Thanyalak Fansiri, Igor Filipovic, Albin Fontaine, Andrea Gloria-Soria, Richard Hall, Vinita S. Joardar, Andrew K. Jones, Raissa G. G. Kay, Vamsi K. Kodali, Joyce Lee, Gareth J. Lycett, Sara N. Mitchell, Jill Muehling, Michael R. Murphy, Arina D. Omer, Frederick A. Partridge, Paul Peluso, Aviva Presser Aiden, Vidya Ramasamy, Gordana Rasic, Sourav Roy, Karla Saavedra-Rodriguez, Shruti Sharan, Atashi Sharma, Melissa Laird Smith, Joe Turner, Allison M. Weakley, Zhilei Zhao, Omar S. Akbari, William C. Black, Han Cao, Alistair C. Darby, Catherine A. Hill, J. Spencer Johnston, Terence D. Murphy, Alexander S. Raikhel, David B. Sattelle, Igor V. Sharakhov,

Bradley J. White, Li Zhao, Erez Lieberman Aiden, Richard S. Mann, Louis Lambrechts, Jeffrey R. Powell, Maria V. Sharakhova, Zhijian Tu, Hugh M. Robertson, Carolyn S. McBride, Alex R. Hastic, Jonas Korlach, Daniel E. Neafsey, Adam M. Phillippy, and Leslie B. Vosshall. 2018. 'Improved reference genome of *Aedes aegypti* informs arbovirus vector control', *Nature*, 563: 501-07.

Meisel, Richard P., Christopher A. Gonzales, and Hoang Luu. 2017. 'The house fly Y Chromosome is young and minimally differentiated from its ancient X Chromosome partner', *Genome Research*, 27: 1417-26.

Papanicolaou, A., M. F. Schetelig, P. Arensburger, P. W. Atkinson, J. B. Benoit, K. Bourtzis, P. Castanera, J. P. Cavanaugh, H. Chao, C. Childers, I. Curril, H. Dinh, H. Doddapaneni, A. Dolan, S. Dugan, M. Friedrich, G. Gasperi, S. Geib, G. Georgakilas, R. A. Gibbs, S. D. Giers, L. M. Gomulski, M. Gonzalez-Guzman, A. Guillem-Amat, Y. Han, A. G. Hatzigeorgiou, P. Hernandez-Crespo, D. S. T. Hughes, J. W. Jones, D. Karagkouni, P. Koskinioti, S. L. Lee, A. R. Malacrida, M. Manni, K. Mathiopoulos, A. Meccariello, S. C. Murali, T. D. Murphy, D. M. Muzny, G. Oberhofer, F. Ortego, M. D. Paraskevopoulou, M. Poelchau, J. X. Qu, M. Reczko, H. M. Robertson, A. J. Rosendale, A. E. Rosselot, G. Saccone, M. Salvemini, G. Savini, P. Schreiner, F. Scolari, P. Siciliano, S. B. Sim, G. Tsiamis, E. Urena, I. S. Vlachos, J. H. Werren, E. A. Wimmer, K. C. Worley, A. Zacharopoulou, S. Richards, and A. M. Handler. 2016. 'The whole genome sequence of the Mediterranean fruit fly, *Ceratitis capitata* (Wiedemann), reveals insights into the biology and adaptive evolution of a highly invasive pest species', *Genome Biology*, 17: 31.

Wang, Y. H., G. Q. Fang, P. H. Xu, B. L. Gao, X. J. Liu, X. W. Qi, G. J. Zhang, S. Cao, Z. H. Li, X. M. Ren, H. R. Wang, Y. H. Cao, R. Pereira, Y. P. Huang, C. Y. Niu, and S. Zhan. 2022. 'Behavioral and genomic divergence between a generalist and a specialist fly', *Cell Reports*, 41: 111654.

Xu, Li, Hong-Bo Jiang, Jie-Ling Yu, Deng Pan, Yong Tao, Quan Lei, Yang Chen, Zhao Liu, and Jin-Jun Wang. 2023. 'Two odorant receptors regulate 1-octen-3-ol induced oviposition behavior in the oriental fruit fly', *Communications Biology*, 6: 176.

Yang, Yang, Hong-Bo Jiang, Chang-Hao Liang, Yun-Peng Ma, Wei Dou, and Jin-Jun Wang. 2022. 'Chromosome-level genome assembly reveals potential epigenetic mechanisms of the thermal tolerance in the oriental fruit fly, *Bactrocera dorsalis*', *International journal of biological macromolecules*, 225: 430-41.

Yang, Yang, Hongfei Li, Changhao Liang, Donghai He, Hang Zhao, Hongbo Jiang, and Jinjun Wang. 2024. 'Neuropeptide signaling systems are involved in regulating thermal tolerance in the oriental fruit fly', *Journal of Integrative Agriculture*.
