## Supplemental Figure S1 to S8, and will be used for the link to the file on the preprint site. for "The assembly of Y chromosome reveals amplification of genes regulating male fertility in *Bactrocera dorsalis*"

**Supplementary Figures**

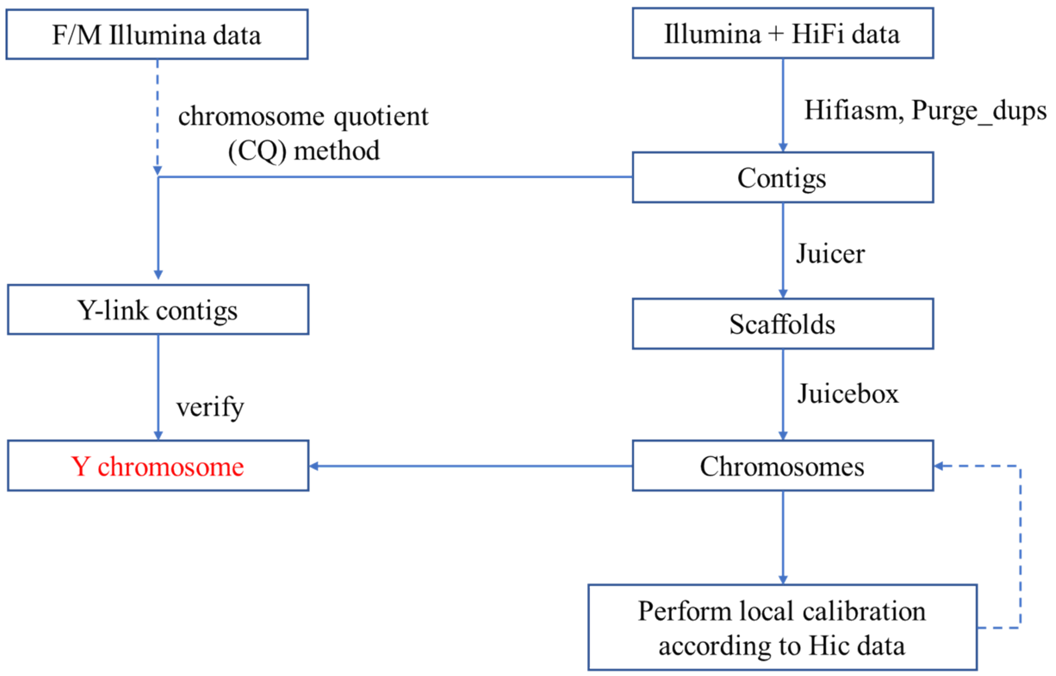


Figure S1. Workflow of genome assembly and Y chromosome identification of *B. dorsalis*.

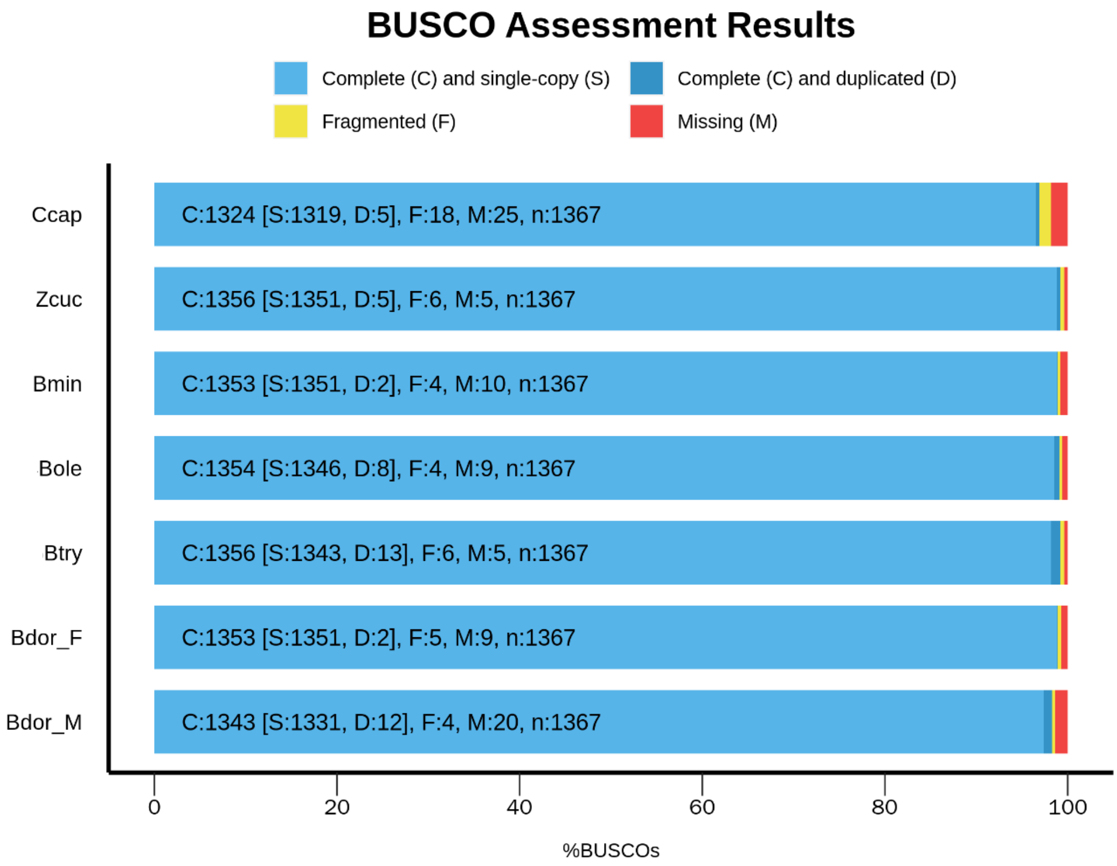


Figure S2. The predicted gene model set completeness of two genders of *B. dorsalis* and 5 other Tephritid flies. The genome completeness and gene set completeness were calculated using BUSCO against insecta_odb10.

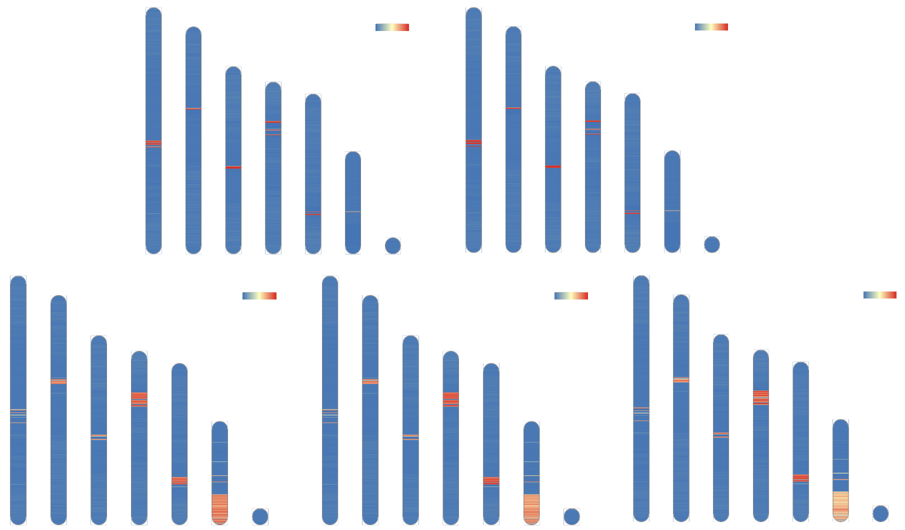

Figure S3. Location of putative centromere-associated repeats. In *B. dorsalis*, five putative centromere-associated repeats were located in the same regions of each chromosome.

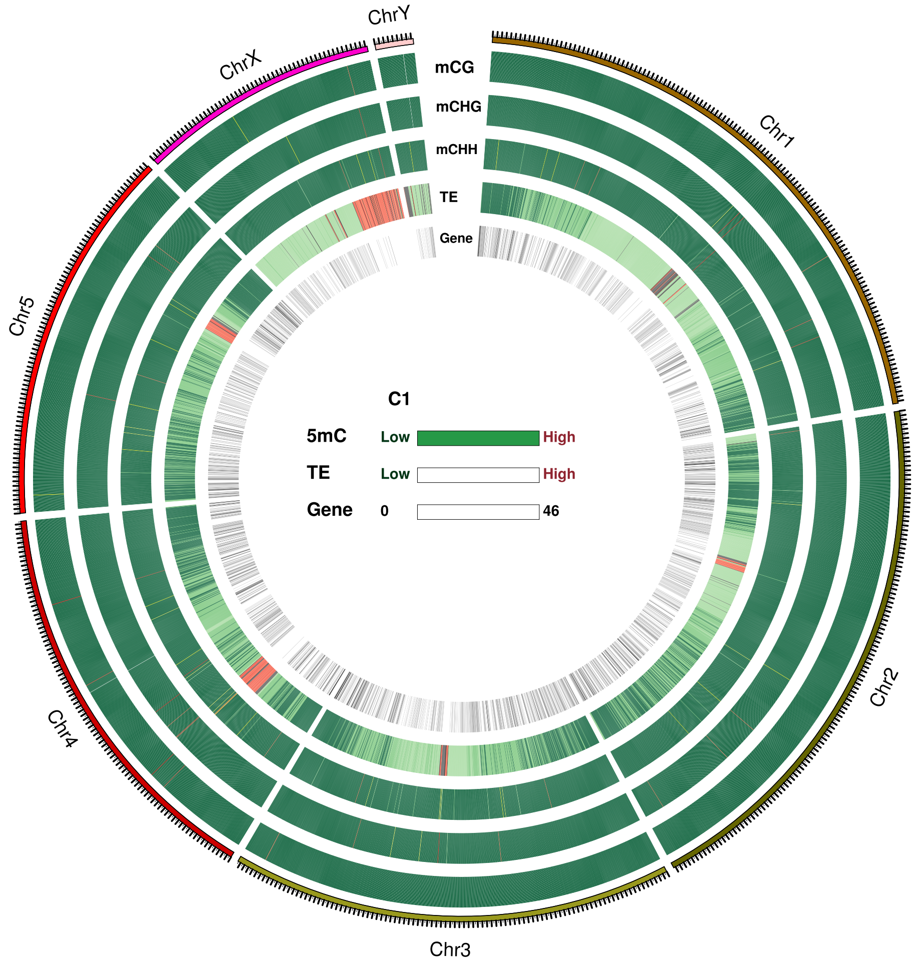


Figure S4. Landscape of methylation, repeat and gene density at chromosome level in *B. dorsalis*. Blocks on the outmost circle represent all seven chromosomes and the denotation of each track are listed in the center of the circle.

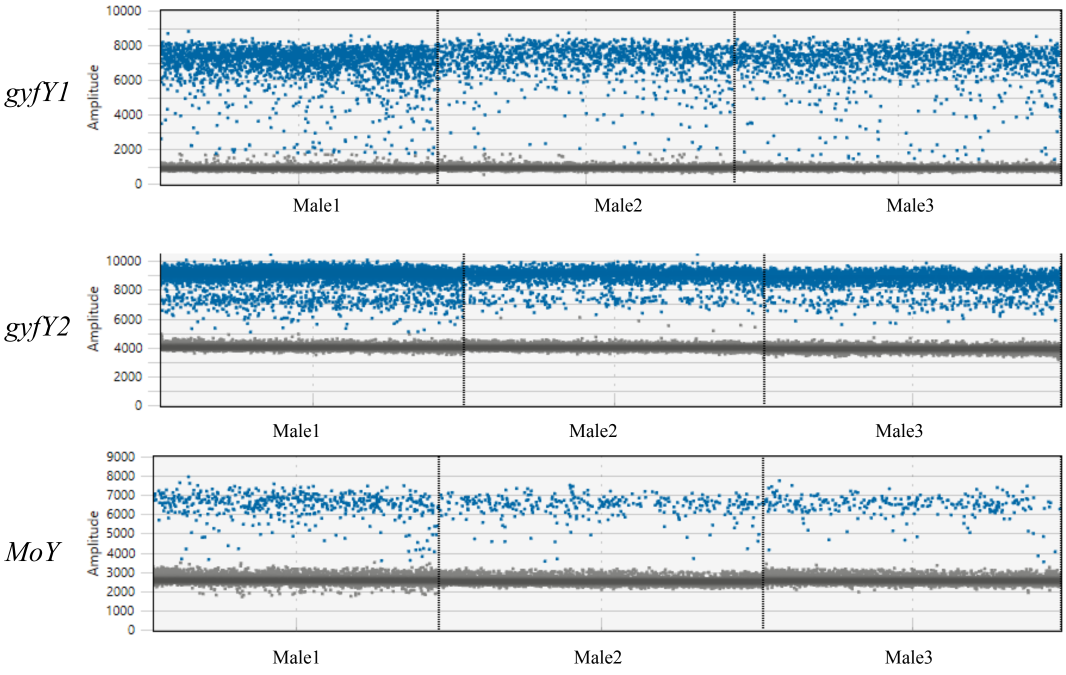


Figure S5. Detection of *gyfY* and *MoY* copy number in male *B. dorsalis* by ddPCR.

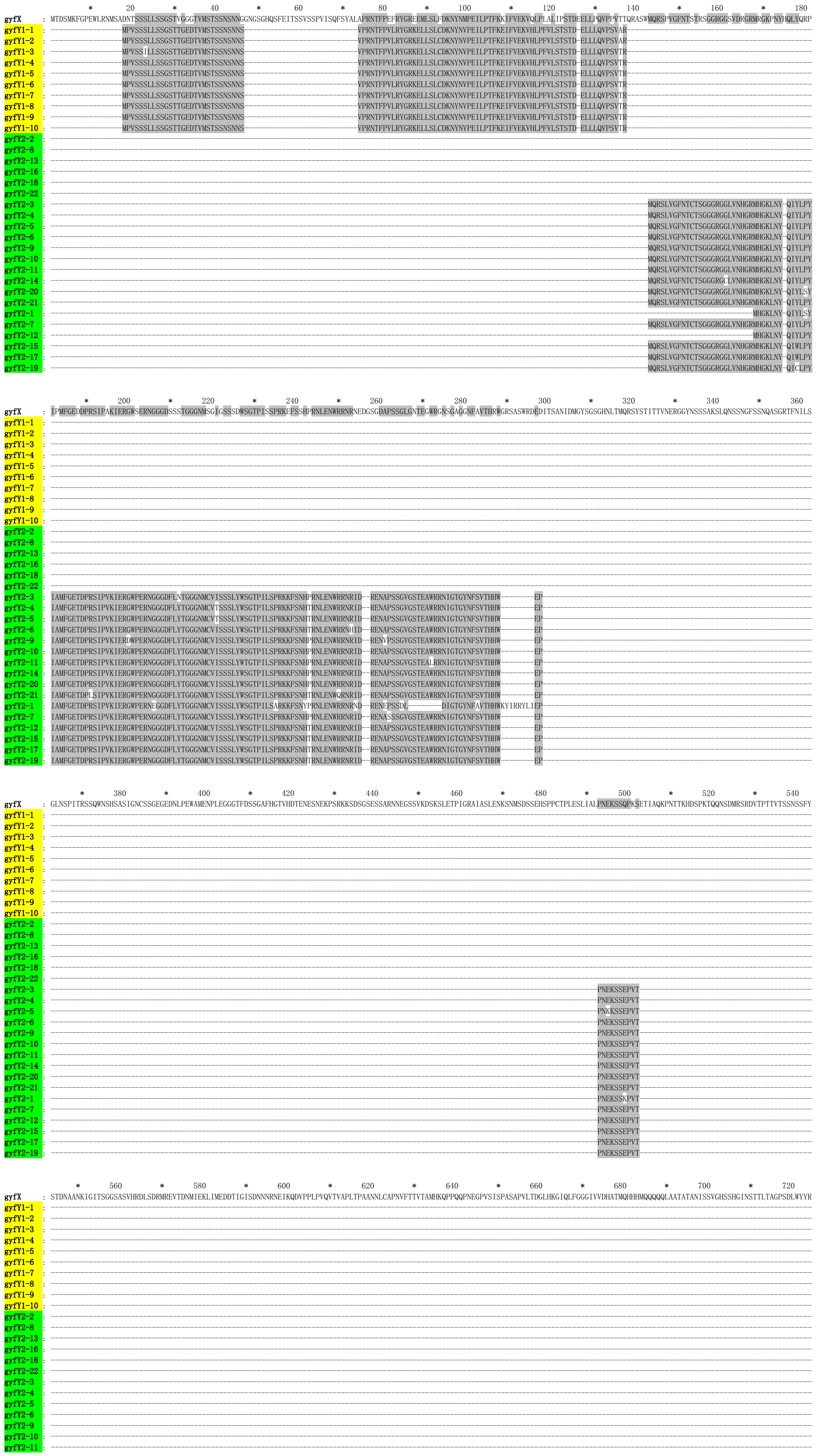

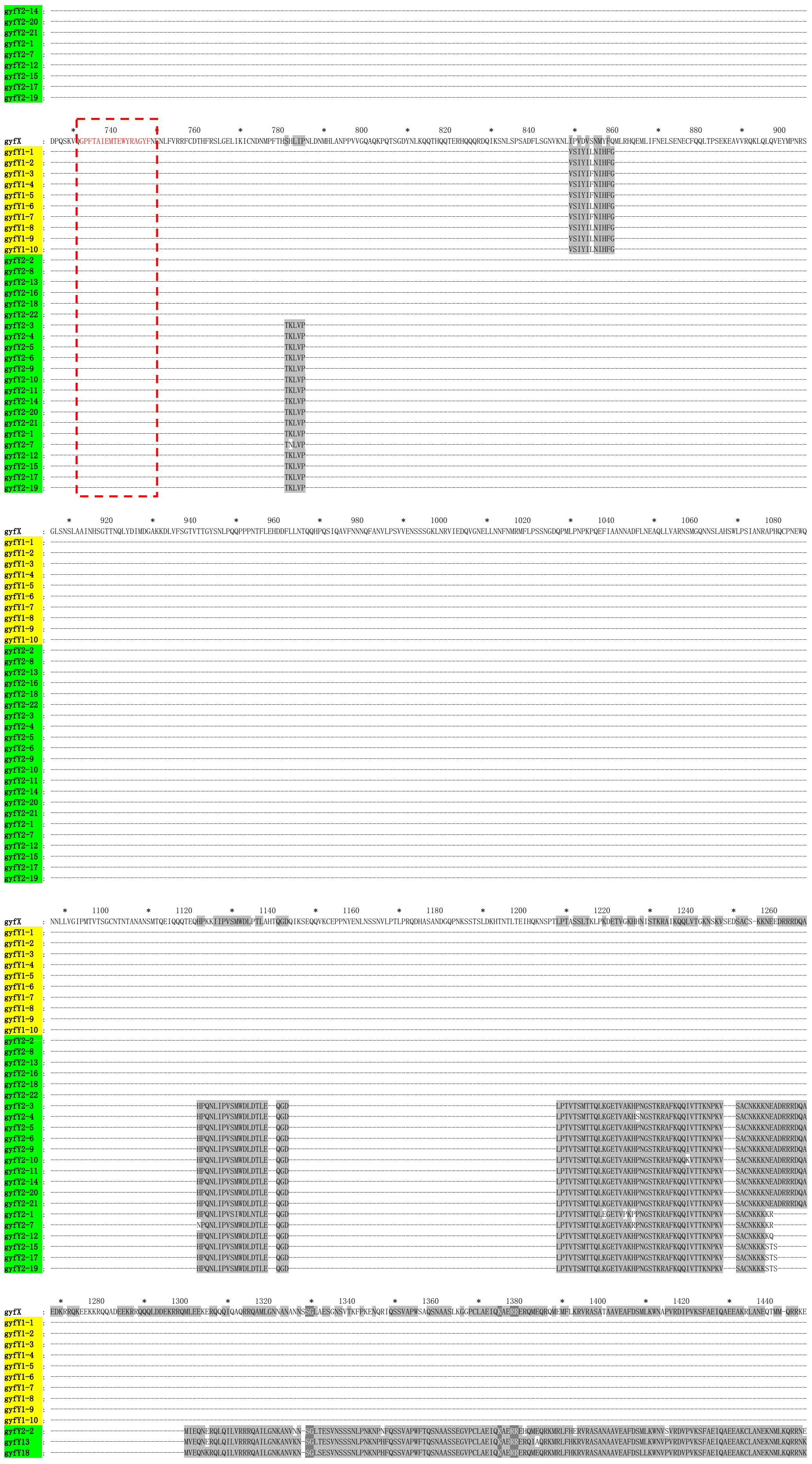
gyfX
gyfY1-1
gyfY1-2
gyfY1-3
gyfY1-4
gyfY1-5
gyfY1-6
gyfY1-7
gyfY1-8
gyfY1-9
gyfY1-10
gyfY2-2
gyfY2-8
gyfY2-13
gyfY2-16
gyfY2-18
gyfY2-22
gyfY2-3
gyfY2-4
gyfY2-5
gyfY2-6
gyfY2-9
gyfY2-10
gyfY2-11
gyfY2-14
gyfY2-20
gyfY2-21
gyfY2-1
gyfY2-7
gyfY2-12
gyfY2-15
gyfY2-17
gyfY2-19
gyfY13
gyfY18

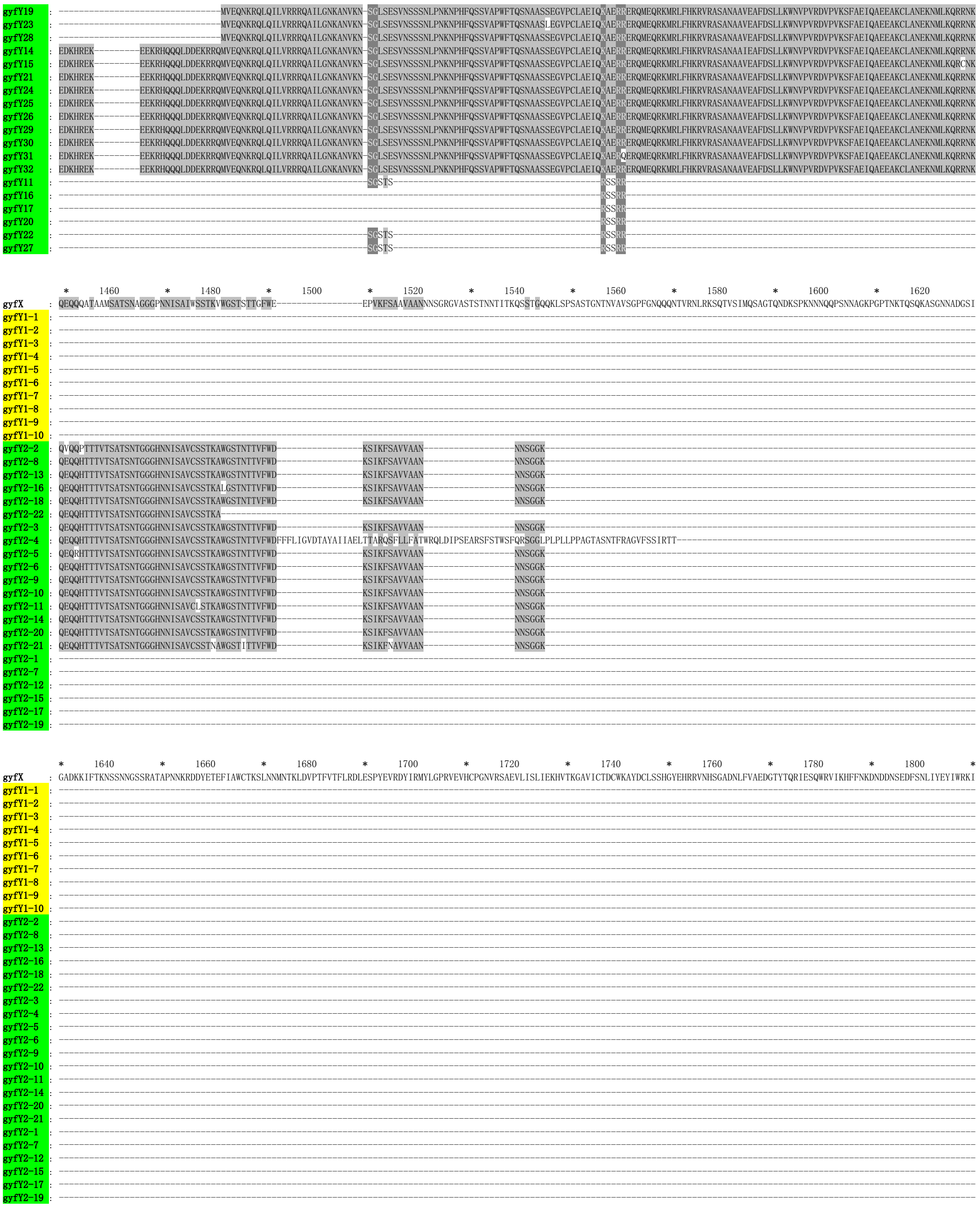

Figure S6 Alignment of *B. dorsalis* proteins gyfY and gyfX. gyfY1 was marked with a yellow background and gyfY2 was marked with a green background. The gyfY has multiple deletions relative to gyfX, including removal of the core GYF motif (red boxed).

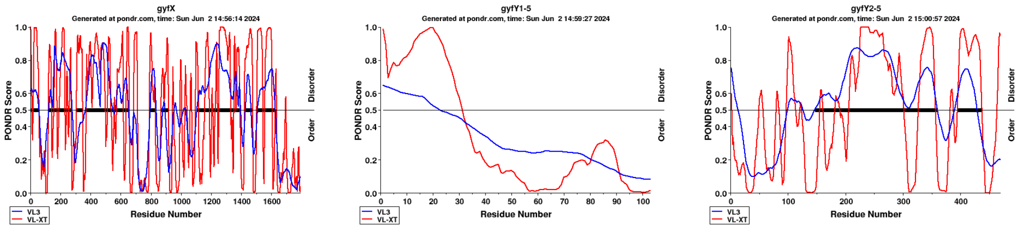


Figure S7. Predicted protein disorder in gyfX and gyfY, as obtained from PONDR, using the algorithms VL3 and VL-XT.

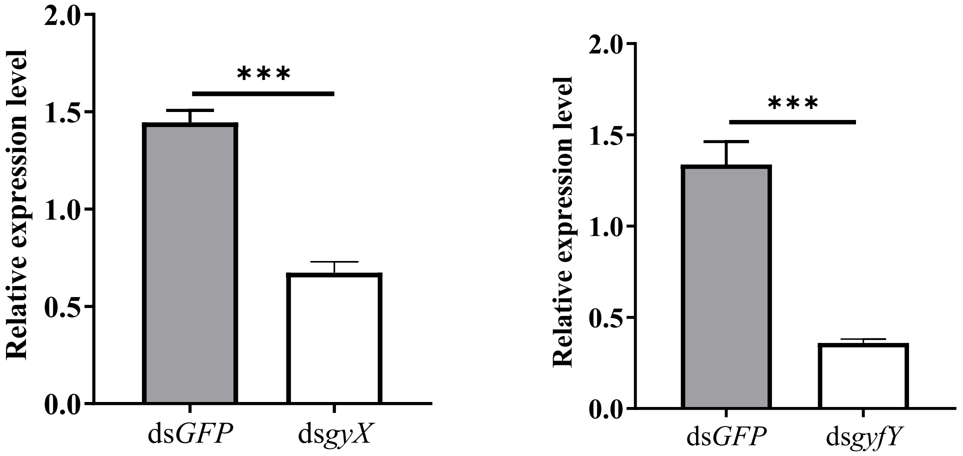


Figure S8. RNAi efficiency was determined by qPCR. Data are means ± standard error. *** indicates $P < 0.001$ (Student's *t*-test) assuming $P = 0.5$ with $n = 4$.
